# Genetic diversity, mobilisation and spread of the yersiniabactin-encoding mobile element ICEKp in *Klebsiella pneumoniae* populations

**DOI:** 10.1101/098178

**Authors:** Margaret M. C. Lam, Ryan R. Wick, Kelly L. Wyres, Claire L. Gorrie, Louise M. Judd, Adam W. J. Jenney, Sylvain Brisse, Kathryn E. Holt

**Affiliations:** Department of Biochemistry and Molecular Biology, Bio21 Molecular Science and Biotechnology Institute, The University of Melbourne, Parkville, Victoria, Australia; Department Infectious Diseases and Microbiology Unit, The Alfred Hospital, Melbourne, Victoria, Australia; Institut Pasteur, Biodiversity and Epidemiology of Bacterial Pathogens, Paris, France

## Abstract

Mobile genetic elements (MGEs) that frequently transfer within and between bacterial species play a critical role in bacterial evolution, and often carry key accessory genes that associate with a bacteria’s ability to cause disease. MGEs carrying antimicrobial resistance (AMR) and/or virulence determinants are common in opportunistic pathogen *Klebsiella pneumoniae*, which are a leading cause of highly drug-resistant infections in hospitals. Well-characterised virulence determinants in *K. pneumoniae* include the polyketide synthesis loci *ybt* and *clb* (also known as *pks*), encoding the iron-scavenging siderophore yersiniabactin and genotoxin colibactin respectively. These loci are located within an MGE called ICE*Kp*, which is the most common virulence-associated MGE of *K. pneumoniae,* providing a mechanism for these virulence factors to spread within the population.

Here we apply population genomics to investigate the prevalence, evolution and mobility of *ybt* and *clb* in *K. pneumoniae* populations through comparative analysis of 2,498 whole genome sequences. The *ybt* locus was detected in 40% of *K. pneumoniae* genomes, particularly amongst those associated with invasive infections. We identified 17 distinct *ybt* lineages and 3 *clb* lineages, each associated with one of 14 different structural variants of ICE*Kp*. Comparison with the wider Enterobacteriaceae population showed occasional ICE*Kp* acquisition by other members. The *clb* locus was present in 14% of all *K. pneumoniae* and 38.4% of *ybt*+ genomes. Hundreds of independent ICE*Kp* integration events were detected affecting hundreds of phylogenetically distinct *K. pneumoniae* lineages, including ≥19 in the globally-disseminated carbapenem-resistant clone CG258. A novel plasmid-encoded form of *ybt* was also identified, representing a new mechanism for *ybt* dispersal in *K. pneumoniae* populations. These data show that MGEs carrying *ybt* and *clb* circulate freely in the *K. pneumoniae* population, including among multidrug-resistant strains, and should be considered a target for genomic surveillance along with AMR determinants.

**AUTHOR SUMMARY:** *Klebsiella pneumoniae* infections are becoming increasingly difficult to treat with antibiotics. Some *K. pneumoniae* strains also carry extra genes that allow them to synthesise yersiniabactin, an iron-scavenging molecule, which enhances their ability to cause disease. These genes are located on a genetic element that can easily transfer between strains. Here, we screened 2498 *K. pneumoniae* genome sequences and found substantial diversity in the yersiniabactin genes and the associated genetic elements, including a novel mechanism of transfer, and detected hundreds of distinct yersiniabactin acquisition events between *K. pneumoniae* strains. We show that these yersiniabactin mobile genetic elements are specifically adapted to the *K. pneumoniae* population but also occasionally acquired by other bacterial members belonging to the Enterobacteriaceae family such as *E. coli.* These insights into the movement and genetics of yersiniabactin genes allow tracking of the evolution and spread of yersiniabactin in global *K. pneumoniae* populations and monitoring for acquisition of yersiniabactin in antibiotic-resistant strains.

## INTRODUCTION

Mobile genetic elements (MGEs) including plasmids, transposons and integrative conjugative elements (ICEs) can generate significant genotypic and phenotypic variation within bacterial populations, driving the emergence of niche- or host-adapted lineages or pathotypes (1,2). Despite the risk that acquisition of virulence-associated MGEs can pose to pathogen emergence, few studies have explored the diversity, distribution and dynamics of such MGEs within their host bacterial populations.

ICE*Kp* is an integrative conjugative element (ICE) that mobilises the *ybt* locus, which encodes biosynthesis of the siderophore yersiniabactin and its receptor (3). Yersiniabactin and other siderophore systems are considered key bacterial virulence factors as they provide mechanisms for scavenging iron (an essential nutrient) from host transport proteins, thereby enhancing the ability of bacteria to survive and replicate within the host (4–6). Nearly all *K. pneumoniae* produce the siderophore enterobactin, however its scavenging mechanisms are inhibited by human lipocalin-2 (Lcn2), which has a strong binding affinity for ferric and aferric enterobactin (7) and induces an inflammatory response upon binding (8). Yersiniabactin escapes Lcn2 binding, thus avoiding the inflammatory response and enhancing bacterial growth and dissemination to the spleen, although it does not provide sufficient iron to allow growth in human serum or urine (8–11). Yersiniabactin can also bind other heavy metals besides iron; for example, yersiniabactin expressed by uropathogenic *E. coli* has been shown to bind Cu^2+^, providing protection against copper toxicity and redox-based phagocyte defences (12). Yersiniabactin is by far the most common *K. pneumoniae* high virulence determinant, present in roughly a third of clinical isolates, and is significantly associated with strains isolated from bacteraemia and tissue-invasive infections such as liver abscess, compared to those from non-invasive infections or asymptomatic colonisation (3,13). In contrast the virulence plasmid, which encodes hypermucoidy and the acquired siderophores salmochelin (which modifies enterobactin to escapes Lcn2 binding (14)) and aerobactin (which can scavenge iron from the host blood protein transferrin (15)), is present in less than 5% of *K. pneumoniae* isolates sampled from infections (13).

The *ybt* locus was first described in the *Yersinia* high pathogenicity island (HPI), variants of which have since been reported in other *Enterobacteriaceae* species (16), including *K. pneumoniae* where *ybt* is located within ICE*Kp* (3,17,18). ICE*Kp* is self-transmissible, involving excision (requiring the gene *xis*), formation of an extrachromosomal circular intermediate (requiring the integrase gene *int* and 17 bp direct repeats at both outer ends), mobilization to recipient cells (requiring *virB1, mobB* and *oriT*) and integration at *attO* sites present in four closely-located tRNA-*Asn* copies in the *K. pneumoniae* chromosome (3,19). ICE*Kp* sometimes carries additional virulence determinants, including *iro* (encoding salmochelin synthesis) or *clb* (also known as *pks*, encoding synthesis of the genotoxic polyketide colibactin (3,17,20), which can induce double strand DNA breaks in eukaryotic cells (21)). Therefore, ICE*Kp* represents a prominent virulence element that strongly influences the pathogenicity of *K. pneumoniae* strains. Although it is significantly associated with invasive infections, the diversity of ICE*Kp* structures and their transmission dynamics within the host bacterial population have not yet been characterized. Here we address this important gap in *K. pneumoniae* virulence evolutionary dynamics by using comparative genomics. We investigate *ybt* phylogenetics, ICE*Kp* structure, integration sites and chromosomal genotypes, and use these signals to track the movement of ICE*Kp* in *K. pneumoniae* populations, and explore the relationship of this MGE with its bacterial host.

## RESULTS

### Diversity of the *ybt* locus in *K. pneumoniae*

First we screened for *ybt* genes in 2498 genomes belonging to the *K. pneumoniae* complex (data sources in **Table S1**), and found *ybt* in 39.5% of 2289 *K. pneumoniae*, but only 2/146 *K. variicola* and 0/63 *K. quasipneumoniae*. Prevalence was 40.0% in the carbapenemase-associated *K. pneumoniae* clonal group (CG) 258, 87.8% in the hypervirulent *K. pneumoniae* CG23, and 32.2% in the wider *K. pneumoniae* population. Consistent with previous reports (3,13), amongst human isolates with reliable clinical source information (**Table S1**), the presence of *ybt* was significantly associated with infection isolates (odds ratio (OR)=3.6, p<1×10^-7^), particularly those from invasive infections (OR=28.6 for liver abscess, OR=4.1 for blood isolates; see **Table S2**).

Each of the 11 *ybt* locus genes displayed substantial diversity within the *K. pneumoniae* population (see **Supplementary Text; Fig. S1, Table S3**). We further explored the genetic diversity of *ybt* using phylogenetic and multi-locus sequence typing (MLST) analyses (see **Methods**). *Ybt* locus sequence types (YbSTs), defined by unique combinations of *ybt* gene alleles, were assigned to 842 *ybt+* isolates (**Table S4**). A total of 329 distinct YbSTs (**Table S5**) were identified, which clustered into 17 phylogenetic lineages (referred to hereafter as *ybt* 1, *ybt* 2, etc; see **Figs. 1, S2, S3**); the phylogenetic tree of translated amino acid sequences was concordant with these lineages (see **Fig. S4, Supplementary Text**).

**Figure 1:**
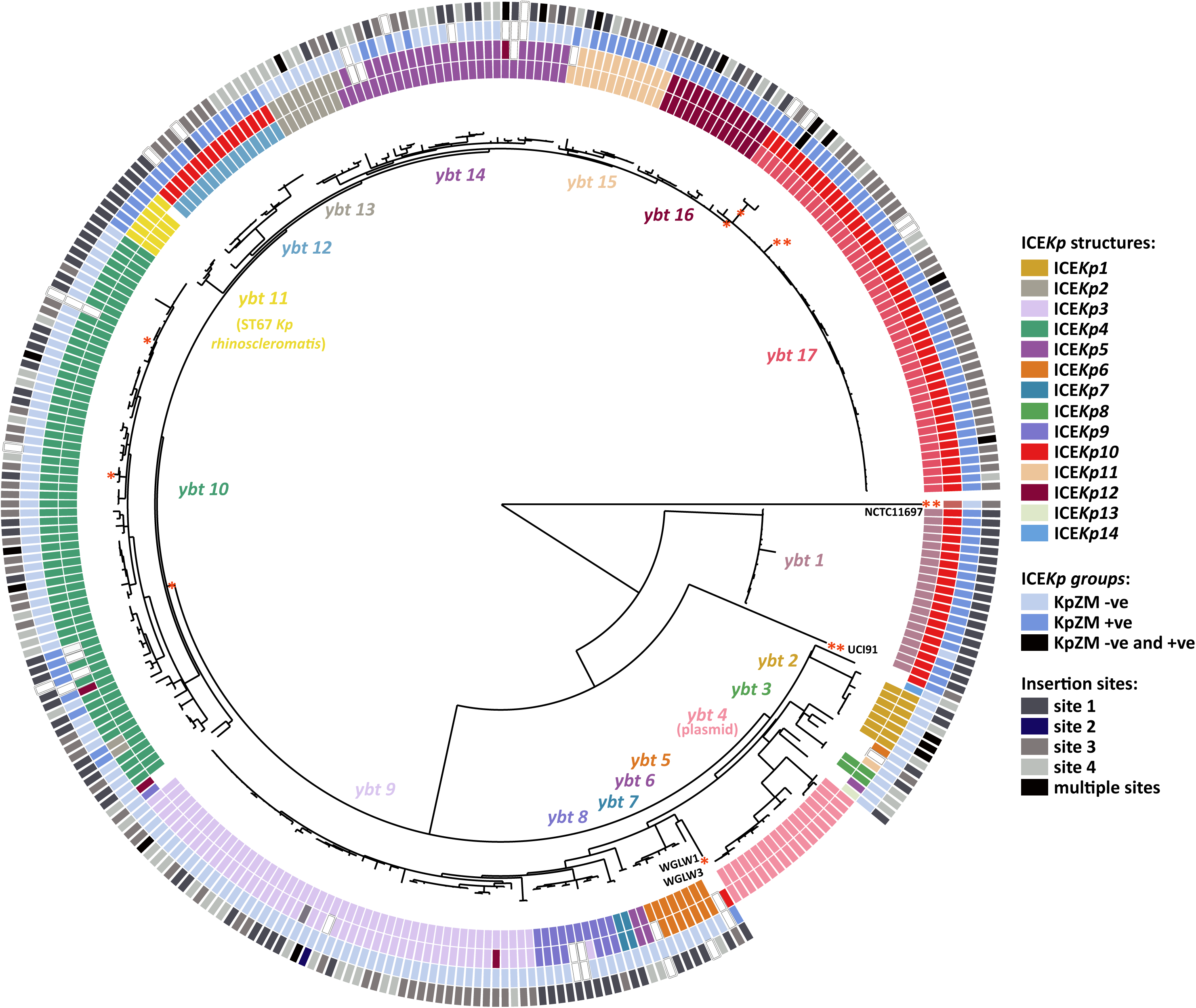
Recombination-filtered phylogenetic analysis of 329 yersiniabactin sequence types identified across 842 genomes. Each leaf represents a single yersiniabactin sequence type (YbST) and these YbST sequences cluster into 17 lineages, as labelled. Tracks (from inner to outer): (1) lineage (key as labelled above the tree nodes; white = unassigned), (2) ICE*Kp* structure (white = undetermined), (3) presence or absence of KpZM module, and (4) tRNA-Asn insertion site (white = undetermined). Recombination events (see **Fig. S2**) are depicted with a red asterisk next to the relevant branches or a single YbST.

Analysis of protein coding sequences dN/dS values was below 0.6 for all genes (**Table S3)**, consistent with moderate purifying selection. Nonsense or frameshift mutations were identified in *ybt* genes in 11% of isolates carrying the *ybt* locus; these mostly affected *irp2* (6.5%) or *irp1* (4%) (see **Table S3)**, which encode key structural components in the yersiniabactin biosynthesis (**Fig. 2A**) (22,23). Inactivation of either of these genes prevents synthesis of yersiniabactin in *Yersinia enterocolitica* (24), and is predicted to have the same effect in *K. pneumoniae*. Most of these mutations (85%) were only observed in a single isolate, suggesting they are not conserved and potentially arose during storage or culture. Consistent with this hypothesis, the presence of inactivating mutations in *ybt* genes was significantly associated with historical isolates (70% amongst *ybt*+ isolates stored since at least 1960, 8% amongst *ybt*+ isolates stored since 2000; OR 27 [95% CI 11-74], p<10^-13^ using Fisher’s exact test). The notable exceptions were ST67 *K. pneumoniae* subspecies *rhinoscleromatis* genomes (conserved frameshift in *irp2*) and ST3 (conserved nonsense mutation in *irp1*), suggesting negative selection in these lineages.

**Figure 2:**
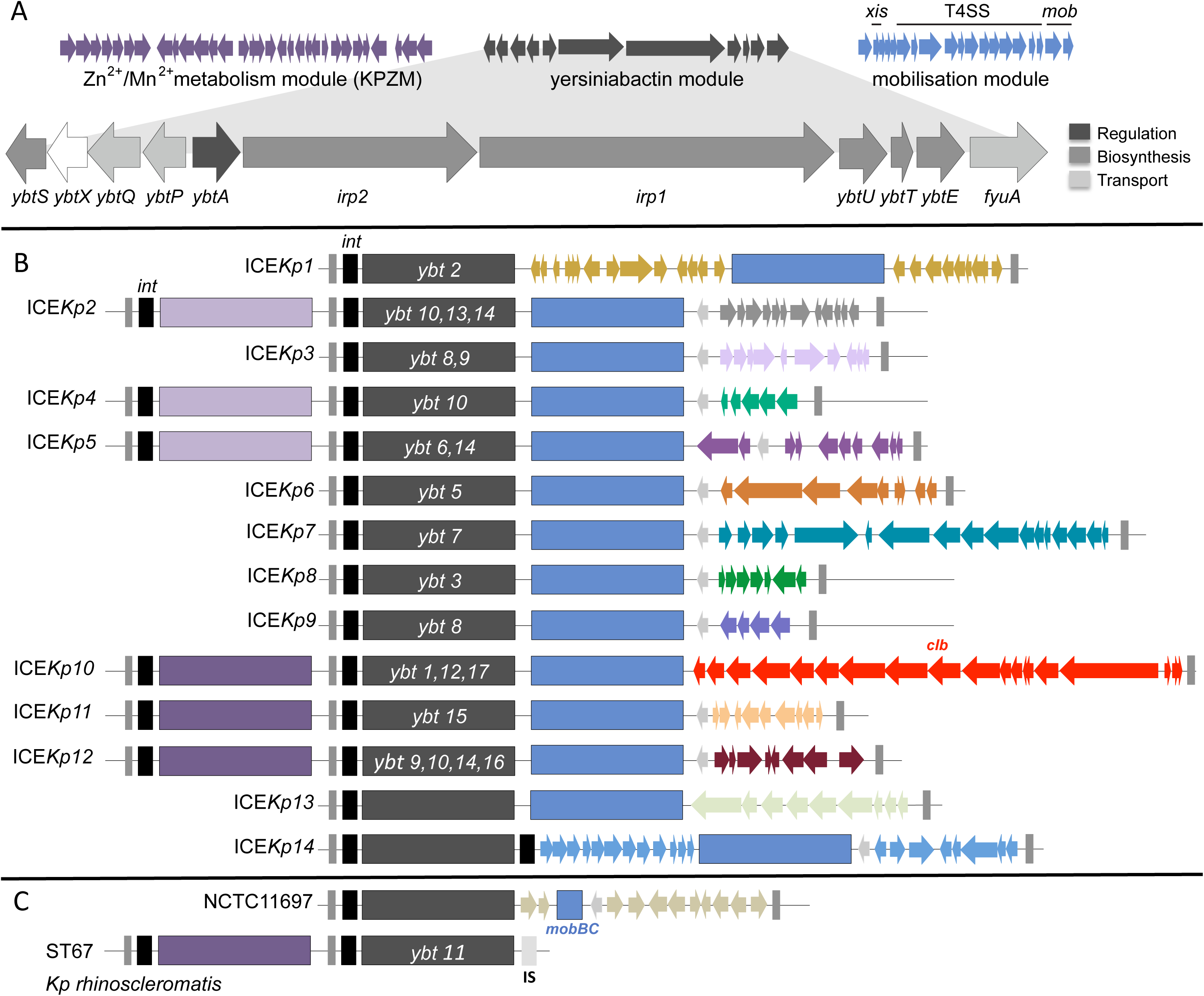
ICE *Kp* structures. (A) Gene structure for core modules, which are shown in **B-C** as coloured blocks: yersiniabactin synthesis locus *ybt* (dark grey, labelled with the most commonly associated *ybt* lineage if one exists), mobilisation module (blue) and Zn^2+^/Mn^2+^ module (purple = usually present, light purple = rarely present). In panels A-B, the variable gene content unique to each ICE*Kp* structure, which is typically separated from the mobilisation module by an antirestriction protein (light grey arrow), is shown in a unique colour as per **Fig. 1**. Grey rectangles represent direct repeats; black rectangles, P4-like integrase genes. Genes that make up the *ybt* locus are also shown and shaded according to their overall role in yersiniabactin synthesis (further details provided in **Table 2**). **(B)** Genetic structures of apparently intact ICE*Kp* variants (see **Table S6** and GenBank deposited sequences for details of specific genes). **(C)** Disrupted ICE*Kp* loci.

### Diversity of ICE *Kp* structures and integration sites in *K. pneumoniae*

With the exception of *ybt* 4 (which we found to be plasmid-borne and representing a novel mechanism of *ybt* transfer in *K. pneumoniae*; see **Supplementary Text**), all *ybt* loci detected in the *K. pneumoniae* genomes were located within an ICE*Kp* structure integrated into one of four copies of tRNA-*Asn* located in a chromosomal region that spans 16.4 kbp in size in strains lacking MGE insertions at these sites (**Fig. S5**). Examples of ICE*Kp* integration were observed at all four tRNA-*Asn* sites (**Fig. 1**), but the frequency of integration differed substantially by site: 35.7%, 44.7%, 19.5% for sites 1, 3 and 4 respectively, and just one integration at site 2. Multiple ICE*Kp* integration sites were observed for most *ybt* lineages (**Fig. 1**); thus, there is no evidence that ICE*Kp* variants target specific tRNA-*Asn* copies.

The boundaries of each ICE*Kp* variant were identified by the 17 bp direct repeats formed upon integration (3), and their structures were compared. This confirmed that ICE*Kp* structures identified in *K. pneumoniae* share several features: (i) a P4-like integrase gene, *int*, at the left end; (ii) the 29 kbp *ybt* locus; (iii) a 14 kbp sequence encoding the *xis* excisionase, *virB*-type 4 secretion system (T4SS), *oriT* transfer origin, and *mobBC* proteins (responsible for mobilisation) (3). In addition, we found that each ICE*Kp* carried a distinct cluster of cargo genes at its right end, which we used to classify the ICEs into 14 distinct structures which we labelled as ICE*Kp2*, ICE*Kp3*, etc, preserving the original nomenclature of ICE*Kp1* (3,25) (see **Fig. 2B**; **Table S6**). We detected occasional additional gene content variation between ICE*Kp* sequences, arising from transposases and other insertion or deletion events. Most of the 14 ICE*Kp* structures were uniquely associated with a monophyletic group in the *ybt* nucleotide-based phylogeny (i.e. a single *ybt* lineage; see **Figs. 1**, **2**; **Table S6**); the exception was the ICE*Kp10* structure, which carries the *clb* locus in the cargo region and was associated with *ybt* lineages 1, 12 and 17 (details below).

All ICE*Kp* carried the integrase gene *int*; however we identified two variant forms of ICE*Kp* lacking the mobilisation genes. First, *K. pneumoniae* subspecies *rhinoscleromatis* (ST67) genomes carried *ybt* 11 but lacked the entire mobilisation module (**Fig. 2C**); as noted above, they also carried nonsense mutations in *irp2*. Second, strain NCTC 11697 carried a highly divergent *ybt* locus (>2% nucleotide divergence from all other *ybt* sequences, marked with ** in **Fig. 1**) and lacked the *virB*-T4SS and *xis* genes (**Fig. 2C**). A ∼34 kbp Zn^2+^ and Mn^2+^ metabolism module (KpZM) was identified upstream of six different ICE*Kp* structures (**Fig. 2**). This module includes an integrase at the left end that shares 97.5% amino acid identity with that of ICE*Kp*, and the same 17 bp direct repeat was found upstream of both integrases and downstream of ICE*Kp*. It is therefore likely that the entire sequence between the outer-most direct repeats (grey bars in **Fig. 2B**) – including the KpZM module, *ybt* locus, mobility and cargo regions *–* can be mobilised together as a single MGE (see **Supplementary Text**).

ICE*Kp1* was the first yersiniabactin ICE reported in *K. pneumoniae* (3,25) and carries a 18 kbp insertion between the *ybt* and mobilisation genes, which our comparative analyses showed to be quite atypical (see **Fig. 2B**). As previously reported, the inserted sequence encodes *iro* and *rmpA* (which upregulates capsule production and is associated with hypermucoid phenotype) and is homologous to a region on the virulence plasmid pLVPK (3). The only other ICE*Kp* structure in which we identified known *K. pneumoniae* virulence determinants was ICE*Kp10*, whose cargo region harbours the ∼51 kbp colibactin (*clb*) locus. The ICE*Kp10* structure corresponds to the genomic island described in ST23 strain 1084 as GM1-GM3 of genomic island KPHPI208, and in ST66 strain Kp52.145 as an ICE-*Kp1*-like region (17,18). We detected ICE*Kp10* in 40% of ST258, 77% of ST23 and 4.0% of other *K. pneumoniae* genomes including 25 other STs (total 13.85% and 38.43% of all and *ybt+ K. pneumoniae* respectively). Notably, all but three of the ICE*Kp10* strains carried the KpZM module at the left end, suggesting that the *clb* locus (**Table S7**) is usually mobilized within a larger structure including KpZM. MLST analysis of *clb* genes identified 65 CbSTs (**Table S8**), similar to the number of YbSTs detected in ICE*Kp10* (n=86). Phylogenetic analysis of the *clb* locus (excluding *clbJ* and *clbK* (due to a 4153 bp deletion that commonly spans the two genes; see **Fig. S6**, **Supplementary Text**), revealed three *clb* lineages that were each associated with a phylogenetically distinct *ybt* lineage: *clb* 1 (*ybt* 12), *clb* 2A (*ybt* 1) and *clb* 2B (*ybt* 17), consistent with multiple acquisitions of *clb* into the ICE (see **Figs. 1, 3** and **Supplementary Text**).

**Figure 3:**
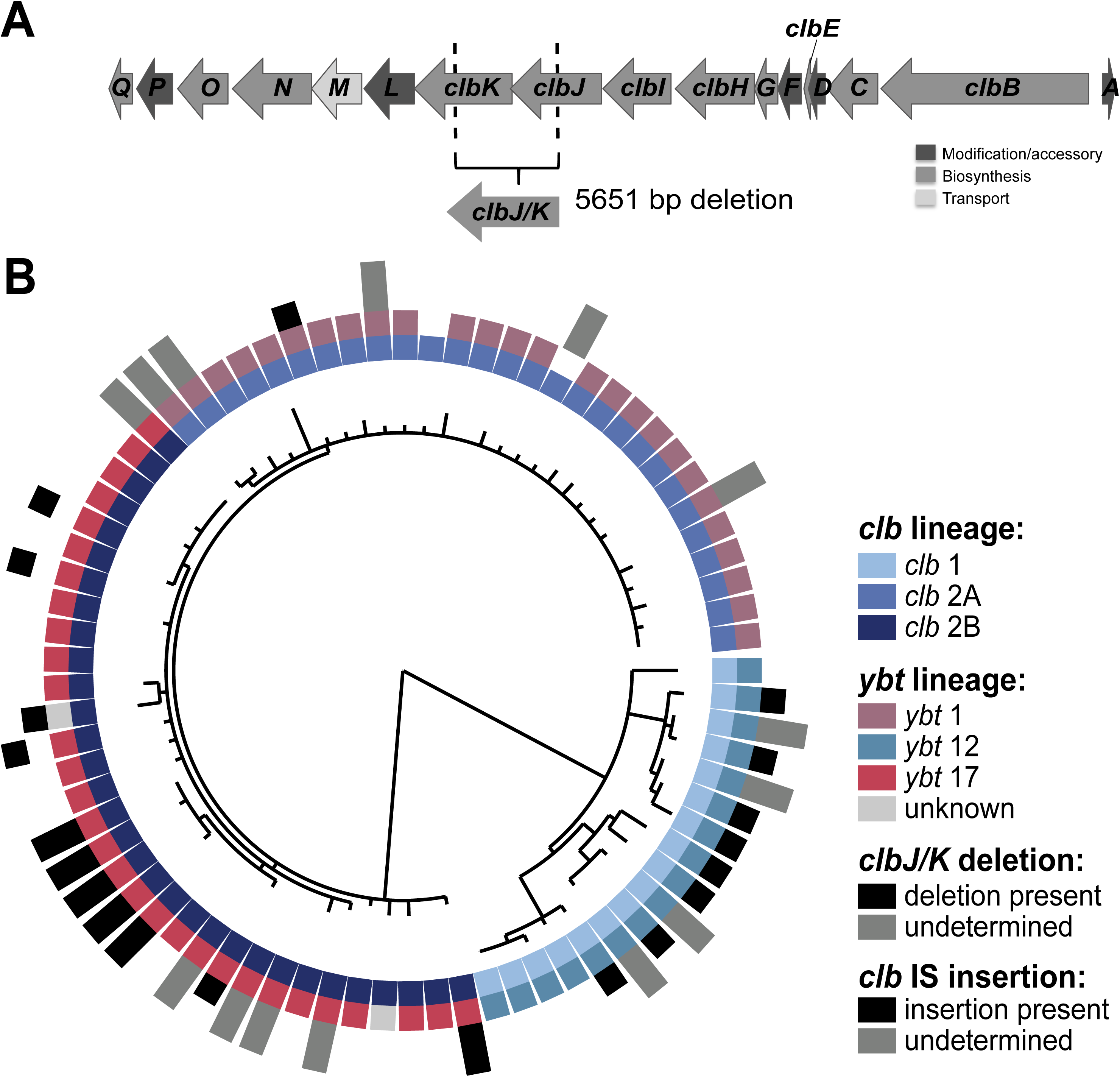
Colibactin diversity. (A) *ClbJ/K* deletion. A 5.6 kbp in-frame deletion between the *clbJ* (6501 kbp) and *clbK* (6465 kbp) genes of the colibactin locus observed in a large number of ICE*Kp10* structures. **(B)** Recombination-free phylogeny of genes in the colibactin locus (excluding *clbJ* and *clbK*). Each leaf represents a unique colibactin locus found across 314 isolates. Tracks (from inner to outer): (1) *clb* lineage, (2) *ybt* lineage (grey = not part of any main *ybt* lineage, white = *ybt-*negative), (3) presence of *clbJ/K* deletion (white = no deletion), and (4) occurrence of intragenic transposase insertions within *clb* locus (white = no insertion).

### Transmission of ICE *Kp* in the *K. pneumoniae* host population

We identified n=206 unique combinations of ICE*Kp* structure, chromosomal ST and integration site. These likely represent distinct *ybt* acquisition events, however it is possible that the ICE*Kp* could migrate to another insertion site following initial integration; hence a more conservative estimate for distinct *ybt* acquisition events is the number of unique combinations of ICE*Kp* structure and chromosomal ST, n=189 (ST phylogenetic relationships are given on **Fig. S7**). The most widely distributed variant was ICE*Kp4* (found in 37 chromosomal STs) followed by ICE*Kp10,* ICE*Kp5* and ICE*Kp3* (n=24, 23 and 23 chromosomal STs, respectively). Conversely, ICE*Kp7*, ICE*Kp8* and ICE*Kp13* were found in one *K. pneumoniae* host strain each (ST111, site 1, n=2; ST37, site 4, n=7; ST1393, site 1, n=1; respectively); ICE*Kp14* was found only in *K. variicola* (ST1986, site 4, n=1).

Twenty-six *K. pneumoniae* chromosomal STs showed evidence of multiple insertion sites and/or ICE*Kp* structures, indicative of multiple integrations of ICE*Kp* within the evolutionary history of these clones (**Figs. 4, S7** and **Table S4**). Most unique acquisition events defined by unique combinations of ICE*Kp* structure and chromosomal STs (65%) were identified in a single genome sequence. The frequency of *ybt* carriage and unique *ybt* acquisitions per ST was correlated with the number of genomes observed per ST (R^2^=0.71, p<1×10^-8^ for log-linear relationship; see **Fig. 5C**), suggesting that the discovery of novel integrations within lineages is largely a function of sampling. This implies that ICE*Kp* may frequently be gained and lost from all lineages, and deeper sampling would continue to uncover further acquisitions and losses. Notably, of the 35 clonal groups that were represented by ≥10 genomes, 30 (86%) included at least one ICE*Kp* acquisition (**Figs. 4, S7**). The five other common clonal groups each consisted mostly of isolates from a localised hospital cluster (ST323, Melbourne; ST490, Oxford; ST512, Italy; ST681, Melbourne; ST874, Cambridge); and we predict that more diverse sampling of these clonal groups would detect ICE*Kp* acquisition events.

**Figure 4:**
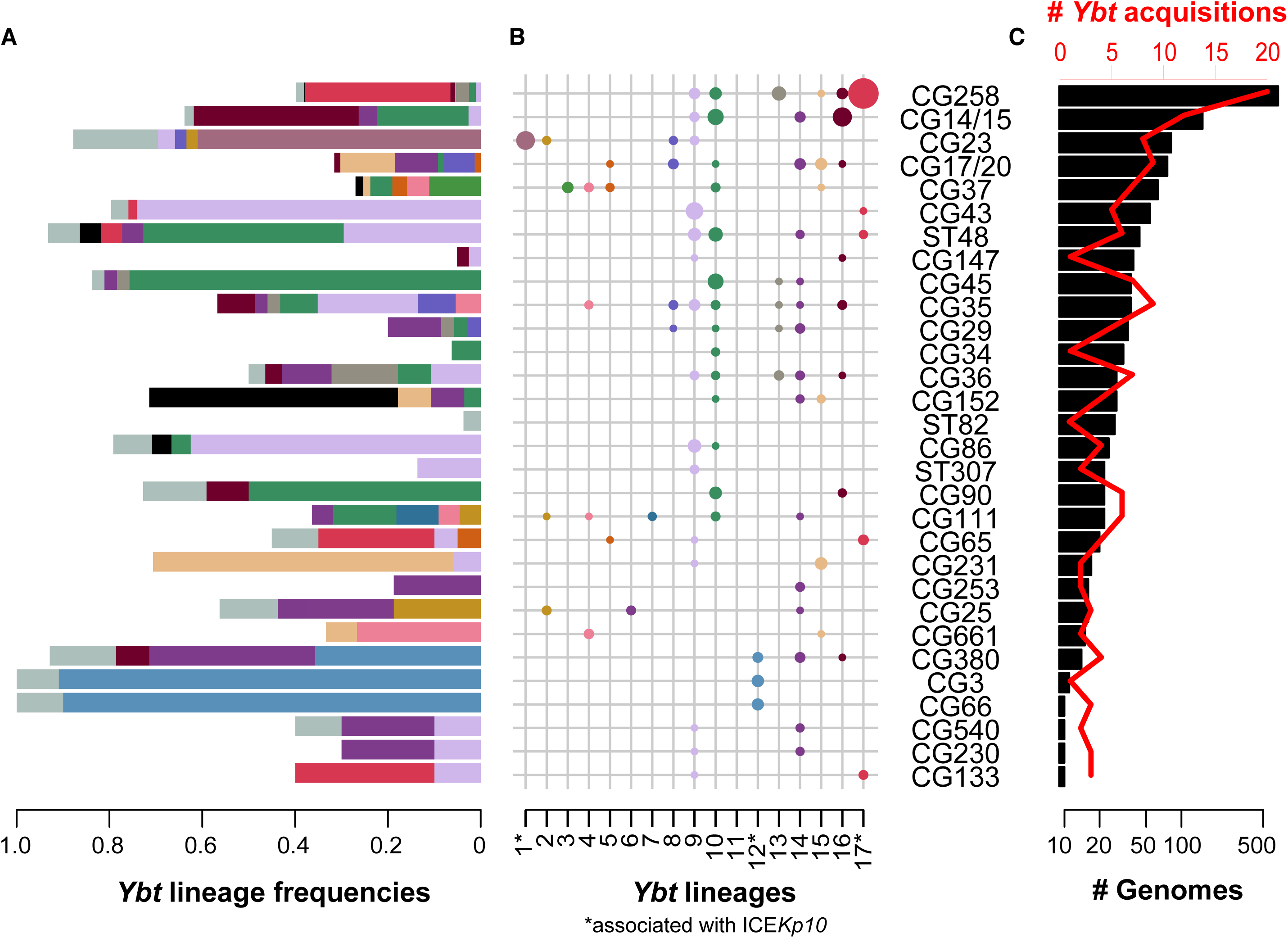
Frequency and diversity of *ybt* sequences and acquisition events in common *K. pneumoniae* clonal groups. This analysis includes all clonal groups for which there were ≥10 genomes available for analysis; each represents a deep branching lineage of the *K. pneumoniae* core genome phylogeny as shown in **Fig. S7**. **(A)** Frequency of *ybt* presence within each clonal group, coloured by *ybt* lineage according to the scheme in **Figs. 1, 3** and panel **B** (white = proportion of genomes without *ybt*). **(B)** Number of *ybt-*positive *K. pneumoniae* genomes designated to a particular *ybt* lineage. The sizes correspond to sample size of genomes. **(C)** Number of genomes (black bars, bottom axis) and number of independent *ybt* acquisition events (red, top axis) per clonal group. Independent *ybt* acquisition events were defined as unique combinations of ICE*Kp* structure and insertion site in each ST.

Of the unique ICE*Kp* acquisition events that were detected in more than one genome, 68% (n=50/73) showed diversity in the YbST. This diversity was attributable to minor allelic changes (SNPs in a median of 2.5 loci), consistent with clonal expansion of ICE*Kp*-positive *K. pneumoniae* strains and diversification of the *ybt* locus *in situ*. The greatest amount of YbST diversity within such groups was observed in hypervirulent clones ST23 (18 YbSTs of ICE*Kp10*/*ybt* 1 in site 1), ST86 (12 YbSTs of ICE*Kp3* in site 3) and ST67 *K. pneumoniae* subspecies *rhinoscleromatis* (five YbSTs in site 1); followed by hospital outbreak-associated MDR clones ST15 (six YbSTs of ICE*Kp4* in site 1 and five in site 3), ST45 (five YbSTs of ICE*Kp4*), ST101 (five YbSTs of ICE*Kp3* in site 3) and ST258 (detailed below). This level of diversity suggests long-term maintenance of the ICE*Kp* in these lineages, allowing time for the *ybt* genes to accumulate mutations.

Given the clinical significance of the carbapenemase-associated CG258 (26), we explored ICE*Kp* acquisition in these genomes in greater detail. *Ybt* was detected in 269 CG258 isolates (40%) from 17 countries; 218 isolates also carried *clb* (nearly all from USA; see **Table S4**). Fifty-eight YbSTs were identified amongst CG258 isolates and clustered into seven *ybt* lineages associated with six ICE*Kp* structures. Comparison of *ybt* lineage, ICE*Kp* structure and insertion site with a recombination-filtered core genome phylogeny for CG258 indicated dozens of independent acquisitions of ICE*Kp* sequence variants in this clonal complex (**Fig. 5**). Near-identical *clb* 2B (ICE*Kp10*/*ybt* 17) sequences were identified in 211 ST258, mostly at tRNA-*Asn* site 3, isolated in the USA during 2003–2014. Most of these isolates carried the *clbJ*/*clbK* deletion (n=175, 83%), and also transposase insertions within other *clb* genes (n=173, see **Table S4**) that may prevent colibactin production (**Fig.5**). A total of 27 *ybt+clb+* ST258 isolates had an apparently intact *clb* locus; two were isolated in Colombia in 2009 and the rest from USA during 2004–2010 (**Table S4**), including the previously reported KPNIH33 (27). The results demonstrate the very high genetic and functional dynamics of ICE*Kp* within a very recently emerged *K. pneumoniae* epidemic lineage, estimated to have emerged in the mid-1990s (28).

**Figure 5:**
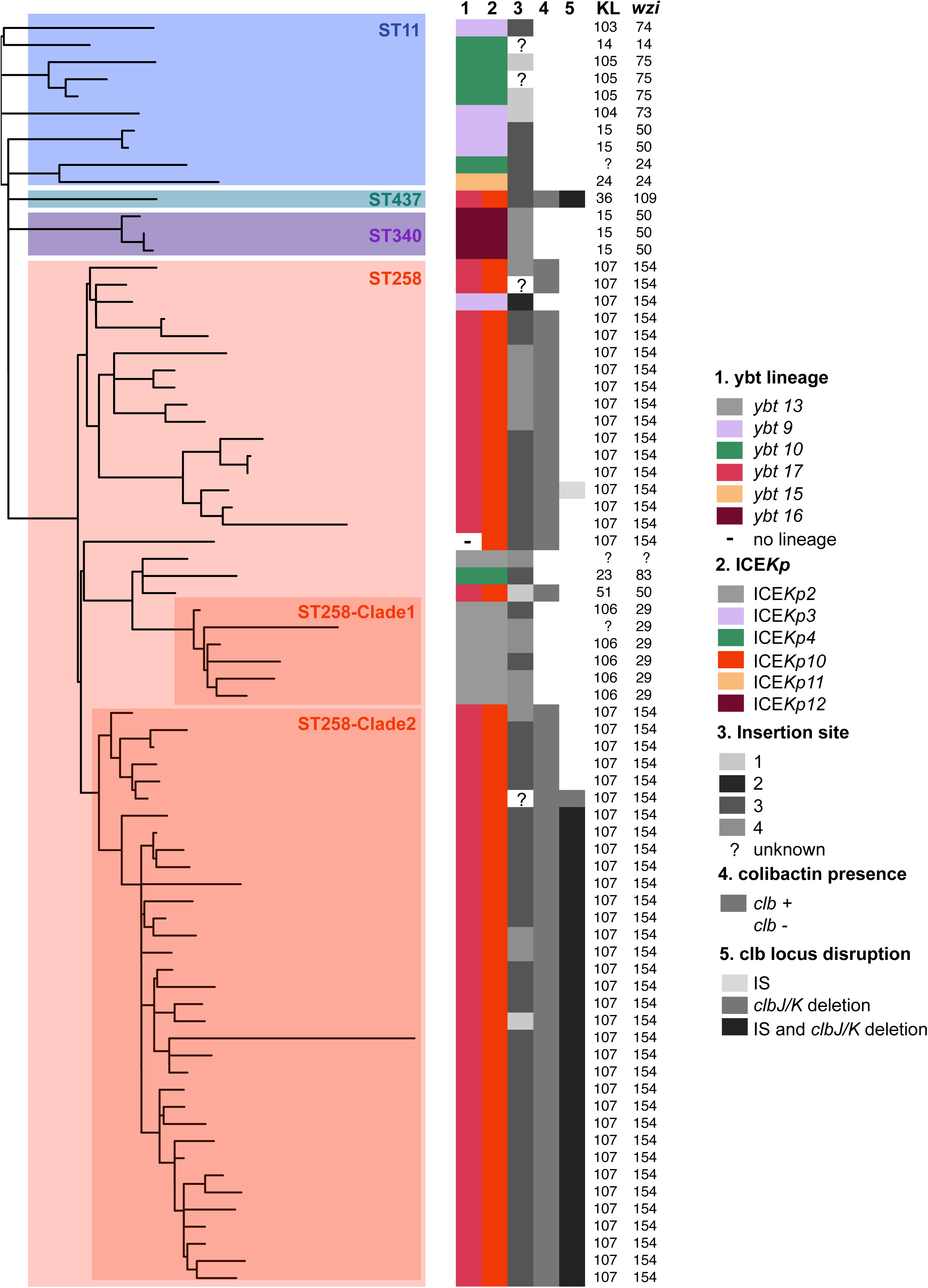
Diversity of *ybt* loci acquired by CG258. The tree shown is an outgroup-rooted maximum likelihood phylogeny inferred from recombination-filtered core gene SNPs for all ybt-positive CG258 isolates. Clades are labelled by chromosomal MLST and the two ST258 clades identified in Deleo *et al.*; tracks indicate (1) *ybt* lineage, (2) ICE*Kp* structure, (3) ICE*Kp* insertion site, (4) *clb* presence/absence and (5) *clb* locus disruptions, coloured according to the inset legend. The K-locus, as determined by Kaptive, and *wzi* allele number are also listed.

### Relationship to *ybt* in other species

Finally, we sought to understand the relationship between ICE*Kp*, which we have shown to circulate within *K. pneumoniae*, and the *ybt* loci found in other bacterial species. BLAST searching NCBI GenBank identified the *ybt* locus in n=242 genomes from 11 species outside the *K. pneumoniae* complex (**Table S9**); all belonged to *Enterobacteriaceae*. The phylogenetic and structural relationships of these loci with those found in *K. pneumoniae* are shown in **Fig. 6**, which indicates that *ybt* sequences mobilised by ICE*Kp* form a subclade that is strongly associated with the species *K. pneumoniae.* The vast majority of ICE*Kp* sequences were found in *K. pneumoniae* (97%); the exceptions were 15 *E. coli*, 6 *Klebsiella aerogenes*, 4 *Citrobacter koseri*, 2 *K. variicola* and 1 *Enterobacter hormaechei* (29); see **Fig. 6, Tables S9**). These include one novel ICE*Kp* variant in *E. coli* strain C8 (accession CP010125.1), however this was also detected in a recently sequenced *K. pneumoniae* draft genome (accession GCF_002248635.1). ICE*Kp* accounted for just 11% of *ybt* sequences detected outside *K. pneumoniae*, and in the remaining 89% was not associated with ICE*Kp* or any other identifiable conjugative machinery. Notably, *ybt* sequences from the HPI of the *Yersinia pseudotuberculosis*/*pestis* complex clustered within the ICE*Kp* clade of *ybt* sequences, and not with the *ybt* sequences from *Yersinia enterocolitica* (**Fig. 6**). This shows that the HPI found in the *Y. pseudotuberculosis* complex is derived from ICE*Kp*. Further, the HPI shares the *int* (**Fig. S8**) and *xis* of ICE*Kp*, facilitating integration and excision of the HPI, but appears to have lost the conjugative machinery that enables its spread between cells from distinct yersiniae lineages, consistent with previous investigations showing the HPI is unable to self-transmit (30,31). Additionally, the *int* in the HPI of *Y. enterocolitica* and *Y. pseudotuberculosis* are unlikely to encode a functional integrase due to frameshift and nonsense mutations.

**Figure 6:**
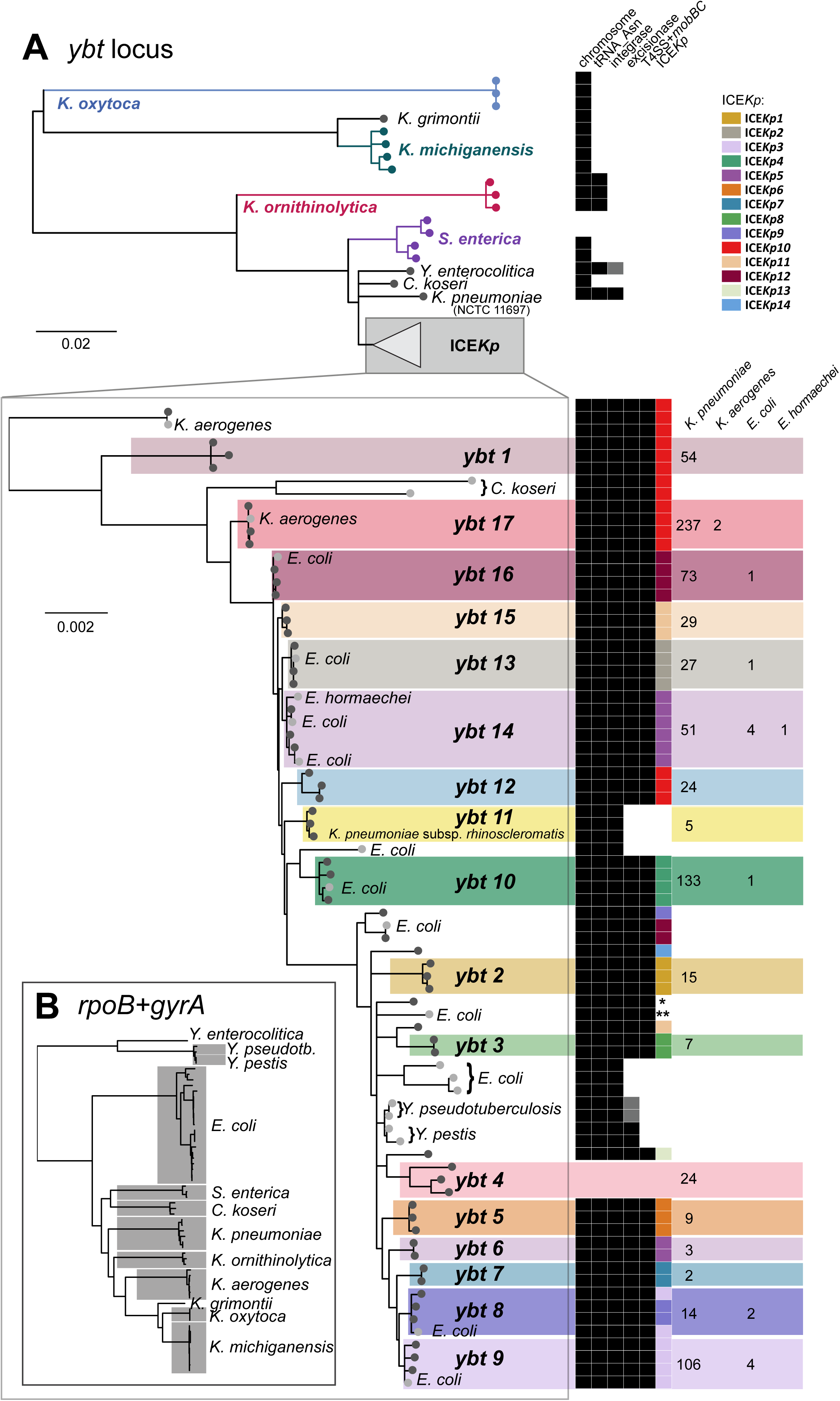
Phylogenetic and structural comparison of *ybt* sequences and corresponding mobile genetic elements in *K. pneumoniae* and other *Enterobacteriaceae* bacteria. (A) Tree shows the midpoint-rooted maximum likelihood phylogeny for DNA sequences of complete *ybt* loci representative of all those found by BLAST search of GenBank, and representatives of the *ybt* lineages identified in this work amongst the *K. pneumoniae* population (shown in **Fig. 1**). Columns on the right are binary indicators of the genetic context of the *ybt* locus in each case: (i) located on the chromosome, (ii) located at a tRNA-*Asn* site, (iii) presence of an integrase (grey indicates non-functional integrase caused by a frameshift or nonsense mutation), excisionase and/or virB-T4SS+*mobBC* portion of the mobility module. For *ybt* sequences in the clade associated with ICE*Kp*, subclades corresponding to the *ybt* lineages (defined in **Fig. 1**) are shaded and additional indicator columns are included to show the corresponding ICE*Kp* structure where present (defined in **Fig. 2**) coloured according to the inset legend (**\*** indicates ICE*Kp* structure not resolvable from available draft genome sequences, **\*\*** indicates novel ICE structure). The number of genomes of each species in which the defined *ybt* lineages was detected is printed on the right. (**B**) Taxonomic relationships between Enterobacteriaceae species in which *ybt* was detected, in the form of a neighbour-joining tree of concatenated *gyrA* and *rpoB* sequences extracted from *ybt*-positive genomes.

The most genetically distant *ybt* loci were found in the chromosomes of *Klebsiella oxytoca* and the related species *Klebsiella michiganensis* and *Klebsiella grimontii* (**Fig. 6**), with no identifiable mobility-associated genes in their proximity; we hypothesise the locus may have originated in the ancestor of these species before becoming mobilised. A related form, situated next to a tRNA-*Asn* but with no identifiable integrase gene, was also present in *Klebsiella ornithinolytica* (also known as *Raoultella ornithinolytica*) (**Fig. 6**). The *ybt* sequences in *Salmonella* formed two related clades, one chromosomal and one plasmid-borne, but lacked proximal mobility-associated genes (**Fig. 6**); notably the plasmid-borne form of *ybt* in *K. pneumoniae* appears to be derived from that of ICE*Kp* and not the plasmid-borne form in *Salmonella* (**Fig. 6**). Finally, this analysis sheds some light on the origins of the atypical form of *ybt* in *K. pneumoniae* NCTC 11697, which includes the integrase but lacks the T4SS mobility region (**Fig 2**), and whose position in the *ybt* tree suggests it may be related to a progenitor element from which ICE*Kp* evolved via acquisition of the excisionase and conjugative machinery.

## DISCUSSION

This study provides key insights into the evolution, dynamics and structure of ICE*Kp*, the most frequent MGE associated with hypervirulent/invasive *K. pneumoniae* infections. The population structure of ICE*Kp* comprises numerous sublineages that are each associated with unique complements of cargo genes in addition to the yersiniabactin synthesis locus *ybt* (**Figs. 1-2**). With the exception of inactivating mutations that likely arise in culture, both *ybt* and *clb* appear to be under strong purifying selection (**Tables S3, S7, Figs. 1 and 3**) and all variants are predicted to synthesise the same yersiniabactin and colibactin polyketide molecules.

The data demonstrate that ICE*Kp* circulates dynamically within the *K. pneumoniae* population. The sheer number of distinct ICE*Kp* acquisitions detected in *K. pneumoniae* (n≥189), and the scale of distinct acquisition events within individual *K. pneumoniae* clonal groups (**Figs. 4-5**), indicates this MGE is highly transmissible within the host bacterial population. Genetic separation of the ICE*Kp* form of *ybt* compared to those in other *Enterobacteriaceae* (**Fig. 6**), and lack of ICE*Kp* outside *K. pneumoniae*, suggests that ICE*Kp* may be specifically adapted to circulate in *K. pneumoniae*. We hypothesise this MGE has been a feature of the *K. pneumoniae* population that predates *K. pneumoniae* sublineage diversification, because (i) the *ybt* genes of ICE*Kp* displayed a similar degree of nucleotide diversity as *K. pneumoniae* core chromosomal genes (mean 0.5%) (13), and (ii) ICE*Kp* was notably rare or absent from *K. pneumoniae*’s closest relatives, *K. variicola* and *K. quasipneumoniae.*

The data indicate that the population prevalence of ICE*Kp* within *K. pneumoniae* (around one third of the population) is sustained through highly dynamic horizontal gene transfer events rather than stable maintenance within *K. pneumoniae* lineages by vertical inheritance. The intermediate frequency of ICE*Kp* in *K. pneumoniae* suggests the existence of some form of balancing selection for the encoded traits. This typically occurs when a trait is most beneficial only when it is not shared by the entire population (e.g. antigenic variation resulting in variable susceptibility to predators or host immunity (32)), or the trait has selective advantages only under certain conditions but high costs in others. Acquisition of *ybt* has benefits in certain iron-depleted conditions, which are presumably encountered in a wide range of environmental and host-associated niches (6); siderophores including yersiniabactin also confer significant growth advantage in heavy metal contaminated soil irrespective of iron content (33). Hence the dynamics of ICE*Kp*/*ybt* may reflect the diverse lifestyles of *K. pneumoniae*, which can vary between hosts and in the environment. However loss of *ybt* also appears to be common, likely occurring due to the high-energy costs from synthesising the polyketide-hybrid molecule. Inactivation of *irp1* or *irp2* in historical isolates (70%) supports strong negative selection against yersiniabactin production in rich media. Notably, *K. pneumoniae* almost universally can synthesise enterobactin, hence the benefit of *ybt* depends on not only the availability of iron but also the form of iron, and other factors such as ability to compete with host iron-binding proteins and evasion of the mammalian immune system’s targeting of enterobactin via Lcn2 (6,8,9). ICE*Kp* cargo genes likely contribute additional costs and benefits to host cells, modifying the fitness equation for their bacterial host; further work will be needed to explore the differential impact and functional relevance of these genes. It is notable that *clb*-carrying ICE*Kp10* was widespread (detected across 32 *K. pneumoniae* lineages) but the *clb* genes were frequently disrupted, suggesting subjection to balancing selection. These disruptions were particularly high amongst the hospital-associated MDR clone ST258, and may indicate selection against costly colibactin production in hospital-adapted strains that already benefit from positive selection under antimicrobial exposure.

Concerningly, our data highlights the possibility that the rate of ICE*Kp* transmission in the population may be sufficiently high that *ybt* is readily available to most *K. pneumoniae* lineages. Hence new clinically important high pathogenicity lineages could theoretically arise at any time following introduction of *ybt* to a strain background that already has features favourable for transmission or pathogenicity in humans, including antimicrobial resistance (AMR). Indeed we found ICE*Kp* to be frequent amongst many of the recognised MDR *K. pneumoniae* clones such as CG258, suggesting that the convergence of AMR and yersiniabactin production is happening frequently in *K. pneumoniae*, potentiating the emergence of lineages that pose substantially greater risk to human health than the broader *K. pneumoniae* population which typically behaves as an opportunistic, mostly susceptible pathogen. FIB_K_ plasmid-borne *ybt* constitutes an entirely novel mechanism for *ybt* mobilisation in *K. pneumoniae*. The FIB_K_ plasmid replicon is very common and highly stable in *K. pneumoniae* but not *E. coli* (34,35), suggesting these plasmids are adapted to *K. pneumoniae* hosts and have the potential to readily transmit *ybt* within the population. Worryingly, many FIB_K_ plasmids have already acquired AMR transposons (36), suggesting there may be few barriers to convergence of AMR and virulence genes in a single FIB_K_ plasmid replicon. Given its potential transmissibility and stability in *K. pneumoniae* hosts, this forms another substantial public health threat and warrants careful monitoring.

The extensive diversity uncovered amongst *ybt* and *clb* sequences and ICE*Kp* structures in this study provides several epidemiological markers with which to track their movements in the *K. pneumoniae* population through analysis of whole genome sequence data, which is increasingly being generated for infection control and AMR surveillance purposes (37,38). The work presented here provides a clear framework for straightforward detection, typing and interpretation of *ybt* and *clb* sequences via the YbST and CbST schemes (**Figs. 1, 3**), which are publicly available in the BIGSdb *K. pneumoniae* database and can be easily interrogated using the BIGSdb web application or using common tools such as BLAST (https://github.com/katholt/Kleborate) or SRST2 (39). In doing so, detection of these key virulence loci provides much-needed insights into the emergence and spread of pathogenic *K. pneumoniae* lineages, which will be particularly important for tracking the convergence of virulence and AMR in this troublesome pathogen.

Of broader relevance, the data show that the deepest diversity of *ybt* sequences is present in the *Klebsiella* genus, and that MGE-borne *ybt* emerged within *K. pneumoniae* before spreading to other *Enterobacteriaceae* (**Fig. 6**). In particular the HPI of *Y. pestis* and *Y. pseudotuberculosis*, where yersiniabactin was first identified and from which it draws its name, is derived from the ICE*Kp* of *K. pneumoniae*; hence the name klebsibactin may have been more appropriate. This adds to the growing body of evidence that *Klebsiella* acts as a reservoir of AMR and pathogenicity genes for other Enterobacteriaceae; KPC and NDM-1 being recent examples of AMR genes first identified in *Klebsiella* that have rapidly become widespread (40,41). We hypothesise this unique role of *Klebsiella* is linked to its more generalist lifestyle, which offers more opportunities to sample accessory genes from a wide array of gene pools. In support of this, *Klebsiella* exhibit extreme differences in gene content within and between species, its accessory genes display a wide range of G+C content and taxonomic sources (13), and strains from environmental niches, such as *K. oxytoca*, can have very large genomes that exceed 6 Mbp. The present work further emphasises the clinical importance of the unique position that *Klebsiella* occupies in the broader microbial sphere as a source of important pathogenicity as well as AMR genes for other Enterobacteriaceae, and should be a motivating factor for further exploration of the ecological and evolutionary mechanisms behind this phenomenon.

## METHODS

### Bacterial genome sequences

We analysed a total of 2498 *K. pneumoniae* genomes (2284 *K. pneumoniae sensu stricto*, 63 *K. quasipneumoniae*, 146 *K. variicola,* 5 undefined or hybrid (13)) obtained from various sources representing a diverse geographical and clonal distribution (**Table S1**; see **Table S4** for full list of isolates and their properties). Just under a third of these genomes had been collected and sequenced in-house from four previous studies of human hospital isolates (13,42–44). These isolates from genotypically and geographically diverse backgrounds, which had clinical source information and were not associated with outbreaks, were used to estimate the distribution of the yersiniabactin locus amongst human isolates associated with the different types of infections listed in **Table S2**.

Where available, Illumina short reads were analysed directly and assembled using SPAdes v3.6.1, storing the assembly graphs for further analysis of genetic context. Where reads were unavailable (n=921), publicly available pre-assembled contigs were used. These had been generated using various strategies and assembly graphs were not available for inspection.

One isolate from our collection (strain INF167, isolated from a patient at the Alfred Hospital, Melbourne, Australia in 2013) was subjected to further sequencing using a MinION Mk1B and R9 Mk1 flow cell (Oxford Nanopore Technologies). A 2D MinION library was generated from 1.5 μg purified genomic DNA using the Nanopore Sequencing Kit (SQK-NSK007). DNA was repaired (NEBNext FFPE RepairMix), prepared for ligation (NEBNextUltra II End-Repair/dA-tailing Module) and ligated with adapters (NEB Blunt/TA Ligase Master Mix). We sequenced the library for 48 hours, yielding 3862 reads (mean length 3049 bp, maximum 44026 bp) that were used to scaffold the SPAdes assembly graph using a novel hybrid assembly algorithm (http://github.com/rrwick/Unicycler). The resulting assembly included one circular plasmid, which was annotated using Prokka (45) and submitted to GenBank under accession KY454639.

### Multi-locus sequence typing (MLST) analysis

Genomes were assigned chromosomal *K. pneumoniae* sequence types by comparison to the 7-locus *K. pneumoniae* MLST scheme (46) in the *K. pneumoniae* BIGSdb database (http://bigsdb.pasteur.fr/klebsiella/klebsiella.html) (47) using SRST2 to analyse reads (39) and BLAST+ to analyse assemblies.

In order to construct novel MLST schemes (48) for the yersiniabactin and colibactin loci, we extracted from the *K. pneumoniae* genome sequences the alleles for genes belonging to the yersiniabactin (*ybtS, ybtX, ybtQ, ybtP, ybtA, irp2, irp1, ybtU, ybtT, ybtE, fyuA*) and colibactin (*clbABCDEFGHIJKLMNOPQR*) synthesis loci, by comparison to known alleles in the *K. pneumoniae* BIGSdb database. To maximise resolution for the novel virulence locus MLST schemes, we included in the definition of sequence types alleles for all 11 genes of the *ybt* locus and 16/18 genes of the *clb* locus (*clbJ* and *clbK* were excluded as they are subject to a common deletion as described in Results). Each observed combination of alleles was assigned a unique yersiniabactin sequence type (YbST, listed in **Table S5**) or colibactin sequence type (CbST, listed in **Table S8**). The schemes and allele sequences are available from the BIGSdb-*K. pneumoniae* website. All genomes with detectable *ybt* or *clb* sequences were included in the definition of YbSTs or CbSTs, with the exception of 61 genomes for which data quality was too low for accurate calling of all alleles (criteria: read depth <20x; <90% agreement of alleles at the read level; and/or incomplete assembly of the *ybt* or *clb* region, which usually was associated with low read depth and generally poor assembly quality with N50 < 100,000 bp).

### Phylogenetic analyses

For each YbST, alignments of the concatenated corresponding allele sequences were produced using Muscle v3.8.31. Recombination events were identified using Gubbins v2.0.0 (49), which screens for regions with a high density of single nuclear polymorphisms (SNPs) that are likely to represent an imported sequence variant. The initial alignment was 28,214 bp long with 2,234 variant sites, of which 232 were identified as recombinant and masked from the alignment (regions shown in **Fig. S2**, visualised using Phandango (https://github.com/jameshadfield/phandango/). Maximum likelihood (ML) trees were inferred from the recombination-masked alignment by running RAxML v7.7.2 (50) five times with the generalised time-reversible (GTR) model and a Gamma distribution, selecting the final tree with the highest likelihood. Lineages were defined as monophyletic groups of YbSTs whose members shared features within the group (≥ 6 shared YbST loci, same ICE*Kp* structures) but were distinguished from other groups (0-1 shared YbST loci, different ICE*Kp* structures). The same approach was used to generate a colibactin ML tree. ML phylogenies for the *ybt* loci and concatenated *rpoB* and *gyrA* sequences from representative *K. pneumoniae* and other *Enterobacteriaceae* bacteria were also generated by running RAxML v7.7.2 (50). Nodes with lower than a bootstrap value of 75 in the *Enterobacteriaceae ybt* phylogeny were collapsed to polytomies with TreeCollapserCL 4 v3.2 (http://emmahodcroft.com/TreeCollapseCL.html).

Core genome SNP trees for *K. pneumoniae* (using one representative genome per each unique ST) and for CG258 (which includes ST258, ST11, ST340 and ST512; using a selection of representative CG258 isolates) were inferred using the mapping pipeline RedDog v1b5 (https://github.com/katholt/reddog) to (i) map short reads against *K. pneumoniae* ST23 strain NTUH-K2044 (25) and ST258 strain NJST258-1 (51), respectively, using Bowtie 2 v2.2.3, and (ii) identify core gene SNPs using SAMtools v1.1. The resulting SNP alignments were subjected to analysis with Gubbins v2.0.0 (to filter recombinant sites), and RAxML v7.7.2 to infer clonal phylogenies.

Translation of the nucleotide sequence into amino acid sequence, inspection of non-synonymous, frameshift and nonsense mutations, and calculations for dN/dS ratios and significance testing for conservation or positive selection were conducted using MEGA 6.06 (52).

### Chromosomal insertion sites and ICE structures

For each *ybt*-positive (*ybt+*) genome, the annotated assembly was manually inspected to determine which of the four tRNA-Asn sites was occupied by ICE*Kp*. This was done with reference to the MGH78578 genome, which lacks any genomic islands at tRNA-Asn sites. The Artemis genome viewer was used to inspect the annotation of the region; BLAST+ was used for genome comparison; and when the region failed to assemble into a single contig, Bandage (53) was used to inspect the locus in the assembly graph where available. Once the insertion site was determined, the structure of the ICE*Kp* was inferred by extracting the sequence between the flanking direct 17 bp repeats ‘CCAGTCAGAGGAGCCAA’, either directly from the contigs using Artemis or from the assembly graph using Bandage. Representative sequences for each ICE*Kp* structure (unless derived from previously assembled genomes) were annotated and deposited in GenBank (accession numbers KY454627 – KY454638).

## Data Availability

Annotated ICE and plasmid sequences generated in this study are available in NCBI Genbank under the accession numbers specified in the text. Yersiniabactin and colibactin MLST schemes are available in the *K. pneumoniae* BIGSdb database at http://bigsdb.pasteur.fr/klebsiella/klebsiella.html. All whole genome sequences analysed in this study are freely available in NCBI, accessions are given in **Table S4**.

## ACKNOWLEDGEMENTS

This work was funded by the NHMRC of Australia (project #1043822 and Fellowship #1061409 to K. E. H) and supported by a Senior Medical Research Fellowship from the Viertel Foundation of Australia (awarded to K. E. H).

## SUPPORTING INFORMATION

**Supplementary text.**

**Supplementary Tables:**

Table S1. Description and sources of genome data used in this study.

Table S2. Frequency of yersiniabactin locus (*ybt*) in *K. pneumoniae* isolated from humans across three previous studies.

Table S3. Summary of function, genetic diversity and mutations observed within *ybt* locus genes.

Table S4. Isolates used in this study (large strain table)

Table S5. Yersiniabactin sequence types (YbSTs) and corresponding alleles. Table S6. Description of ICE*Kp* variants.

Table S7. Summary of function, genetic diversity and mutations observed within *clb* locus genes.

Table S8. Colibactin sequence types (CbSTs) and corresponding alleles Table S9. Description of *ybt* loci detected in *Enterobacteriaceae* bacteria

### Supplementary Figures

**Figure S1. Number and distribution of non-synonymous (ns. mut.) and synonymous mutations (s. mut.) across the translated Irp2 and Irp1 peptides.** The number of mutations are shown on the y-axis of each panel. Non-synonymous and synonymous mutations observed across the *irp2* and *irp1* alleles found in the most divergent sequence of *ybt* are shown on separate plots and labelled accordingly. Key domains within each peptide are highlighted and labelled with domain names and accession numbers.

**Figure S2. Predicted recombination events in the *ybt* locus.**

Recombination events were predicted using Gubbins and are shown as coloured blocks (visualised using Phandango). The number of SNPs introduced (and corresponding % nucleotide divergence) in each recombinant block is indicated. Coordinates along the *ybt* locus and gene boundaries are indicated on the x-axis, with a separate plot showing the total number of recombination events detected. Each row in the plotting area represents a YbST. Phylogenetic relationships between the YbSTs are shown in the tree to the left, which is a midpoint-rooted, recombination-free YbST phylogeny reproduced from **Figure 1**. Colours and numbers on the tree indicate *ybt* lineages as detailed in the text and **Figure 1**.

**Figure S3. Minimum spanning tree of YbSTs, visualised using PhyloViz.** Each node represents a YbST, connections between the nodes indicate allele sharing between YbSTs; nodes are coloured by *ybt* lineages (as defined in **Figure 1**; black indicates no lineage assigned) and labelled with ICE*Kp* structures (as defined in **Figure 2**; the *clb*+ ICEKp10 structure, boxed, is associated with three *ybt* lineages).

**Figure S4. Phylogenetic relationships between the predicted amino acid sequences of 295 YbSTs.**

Each leaf represents a translated amino acid sequence for a yersiniabactin sequence type (YbST), excluding those that encode a frameshift mutation (see **Table S4** for strains with these mutations). The corresponding lineages from **Figure 1** are marked on the outer track and labelled accordingly; white = unassigned.

**Figure S5. Chromosome region in *K. pneumoniae* containing tRNA-Asn sites that are targeted by yersiniabactin ICE *Kp* elements.** The hotspots for insertion of ICE*Kp* and other genomic islands occur within four tRNA-Asn sites, represented by the rectangular blocks, and are marked in the figure. Grey arrows represent coding sequences, labelled by gene symbol or the encoded protein product.

**Figure S6. Conserved domains present in the predicted proteins encoded by (A) *clbJ,* (B) *clbK* and (C) the *clbJ/K* deletion.** The homologous region shared between the amino acid adenylation domains of *clbJ* and *clbK* is shown. The left hand side and a large portion of the fusion product matches to *clbJ* (dark blue) while the right hand side matches to *clbK* (light blue).

**Figure S7. Distribution of *ybt* lineages amongst *K. pneumoniae* chromosomal lineages.** The tree shown is an outgroup-rooted maximum likelihood phylogeny inferred from recombination-filtered *K. pneumoniae* core gene SNPs. Each tip represents a unique *K. pneumoniae* chromosomal ST; clonal groups with ≥10 genome sequences available for analysis are highlighted and labelled. Strain NCTC11697, which carries the most divergent *ybt* sequence, is also highlighted and labelled. The heatmap indicates which *ybt* lineages were detected within each *K. pneumoniae* ST (the relative abundance of *ybt* lineages within each of the highlighted clonal groups are shown in **Figure 4**).

**Figure S8. Phylogenetic relationships between the *int* found in ICE *Kp* and yersiniae HPI.** The tree shown is a neighbour-joining tree of the integrase gene detected in *K. pneumoniae, K. aerogenes, C. koseri* and *E. coli*, and from the HPI in *Y. pseudotuberculosis, Y. pestis* and *Y. enterocolitica.*

